# *Acanthella acuta* Mitochondrial DNA: Linear Architecture and Other Unique Features

**DOI:** 10.1101/2024.01.10.575101

**Authors:** Momin Ahmed, Ehsan Kayal, Dennis V. Lavrov

## Abstract

While *Acanthella acuta* (Schmidt 1862), a common demosponge found in the Mediterranean Sea and Atlantic Ocean, is — morphologically — little distinguishable from other sponges, its mitochondrial DNA (mtDNA) is unique within the class. In contrast to all other studied demosponges, *A. acuta* mtDNA appears to be linear and displays several unusual features: inverted terminal repeats, group II introns in three mt-genes, and two unique ORFs. One of the ORFs (*ORF1535*) combines a DNA-polymerase domain with a DNA-directed RNA-polymerase domain, while the second bears no discernable similarity to any reported sequences. The group II intron within the *cox2* gene is the first such intron reported in an animal. Our phylogenetic analyses indicate that the *cox1* intron is related to similar introns found in other demosponges, while the *cox2* intron is likely not of animal origin. The two domains found within *ORF1535* do not share a common origin and, along with the *cox2* intron, were likely acquired by horizontal transfer. The findings of this paper open new avenues of exploration in the understanding of mtDNA linearization within Metazoa.

**Significance:** The transition from circular to linear mtDNA architecture is a rare and poorly understood phenomenon in animals. In addition, the few lineages that made this transition (Medusozoa, Calcarea, and Isopoda), adopted linear mt-DNA architecture early in their history, making it difficult to reconstruct the underlying events. The identification of linear mtDNA in *Acanthella acuta* not only provides an independent example of the origin of linear mtDNA architectures in animals but also an opportunity to investigate its evolution at a much shorter evolutionary time. In particular, the conservation of circular mtDNA in other studied species within the family Dictyonellidae highlights the need to investigate other species in the genus *Acanthella*, to better understand mtDNA linearization in *A. acuta*.

## Introduction

When the first animal mitochondrial DNAs (mtDNA) were sequenced in the 1980s from human (Anderson et al. 1981) and *Drosophila* (Clary and Wolstenholme 1984), their high degree of synteny and remarkably similar size, shape, and gene content gave rise to the notion of a “typical” metazoan mtDNA. This typical metazoan mtDNA is generally depicted as a small, circular molecule containing a well-conserved set of genes involved in oxidative phosphorylation (OXPHOS) and protein (Gray, Burger, and Lang 1999; Gualberto et al. 2014; Smith and Keeling 2015). While most animal mtDNA sequenced to date follow this pattern, many exceptions have been uncovered, especially in non-bilaterian animals, namely members of Cnidaria, Ctenophora, Placozoa, and Porifera (Lavrov and Pett 2016). In particular, independent transitions from a circular to a linear mitochondrial genome architecture occurred in Medusozoa (Cnidaria) and Calcarea (Porifera) (Bridge et al. 1992; Lavrov et al. 2013). Linear mitochondrial genomes have also been reported in multiple species of Isopoda (Arthropoda) (Doublet et al. 2012).

Linear mtDNA in animals was first documented within the clade Medusozoa (Warrior and Gall 1985), which encompasses traditional classes Hydrozoa, Cubozoa, Scyphozoa, and Staurozoa. To date, more than 70 complete or near complete mt-genomes from this group have been determined, all of them having a linear architecture (Kayal et al. 2012). The few cases of medusozoan mtDNA annotated as circular in GenBank are likely annotation errors (Ahuja et al. 2023). All medusozoan mtDNAs harbor inverted terminal repeats (ITRs) that are hypothesized to function as telomeres (Kayal et al. 2012; Smith et al. 2012). Furthermore, the ancestral medusozoan mtDNA is thought to have contained two extra open reading frames (ORFs): one encoding a type B DNA-polymerase (*polB*) and another of unknown function, though these have been lost in the lineage leading to hydroidolinan hydrozoans (Shao et al. 2006; Kayal and Lavrov 2008; Kayal et al. 2012, 2015; Smith et al. 2012). Linearization of the mtDNA in Medusozoa is hypothesized to have involved the incorporation of a linear plasmid containing *polB* and ITRs (Kayal et al. 2012), a mechanism similar to what has been proposed for linear mtDNAs in fungi (Mouhamadou, Barroso, and Labare 2004).

Linear and multipartite mtDNA has also been documented in calcareous sponges (class Calcarea, phylum Porifera), with lineage-specific characteristics associated with each subclass (Lavrov et al. 2013, 2016). In the calcinean *Clathrina clathrus*, the mtDNA is organized into six linear chromosomes, where up to half of each chromosome is noncoding, while one end of the chromosome is composed of a hairpin-forming noncoding sequence, and the other end comprises a noncoding sequence that is highly similar (61%-98%) across all chromosomes (Lavrov et al. 2013). In calcaroneans, the mt-genome is composed of multiple linear chromosomes. Many of these harbor one or more genes, while some “empty” chromosomes display no discernable coding region (Lavrov et al. 2016).

Among bilaterian animals, linear mtDNA is rare but has been reported within the order Isopoda (subphylum Crustacea). Numerous species from across the order possess a mitogenome consisting of two molecules, a 14 kbp linear monomer and a 28 kbp circular dimer formed from the fusion of two monomers in opposite (“head-to-head”) polarities. This “atypical” mitogenomic organization is patchily distributed across isopods and can be found in both marine and terrestrial species. It is unclear so far if this “atypical” mitogenome organization has convergently evolved multiple times across the different lineages of isopods, or if this represents the ancestral state of isopod mtDNA, which has subsequently been lost in at least three lineages. The presence of the atypical mitogenome organization suggests that the latter might be more likely (Doublet et al. 2012; Peccoud et al. 2017; Pearman et al. 2022)

The transitions described above occurred early in the evolutionary history of those lineages and, among the nonbilaterians, have been conserved among all studied representatives of that group (i.e., Medusozoa and Calcarea). Here we posit that a much more recent transition from circular to linear organization occurred in the demosponge *Acanthella acuta* (Schmidt 1862). Within Porifera, the class Demospongiae is the largest and most diverse. To date, all sampled mtDNA from demosponges have been circular. *Acanthella acuta* is a relatively common demosponge in the Mediterranean and Atlantic oceans. Unlike other closely related species, the mtDNA of *A. acuta* presented several challenges for long-PCR amplification and sequencing. Here we used low-coverage total DNA sequencing to assemble the complete mitochondrial genome for this species and describe unusual features absent from close relatives, including ITRs, ORFs, and multiple introns. We suggest that at least some of these features are directly related to its linear organization.

## Results

### The Acanthella acuta mitogenome

The mtDNA of *A. acuta* has been reconstructed as a linear molecule of >37,199 bp. The overall nucleotide composition was 58.43% A+T with protein-coding genes having the highest %AT (66.30%) and tRNA genes the lowest (55.18%). Table 1 shows a breakdown of %AT content along with AT and GC skews.

**Table 1.** The nucleotide content and skews by different regions of the A. acuta mitogenome.

| Region | %AT | AT Skew | GC Skew |
| --- | --- | --- | --- |
| Protein coding | 66.30 | -0.107 | 0.104 |
| tRNA | 55.18 | -0.008 | 0.103 |
| rRNA | 58.06 | 0.149 | 0.098 |
| Intergenic | 56.58 | 0.036 | 0.006 |

We identified the typical set of 14 poriferan mitochondrial protein genes coding for ATP synthase subunits 6, 8, and 9 (*atp6, 8, 9*), cytochrome oxidase subunits 1-3 (*cox1–3*), NADH dehydrogenase subunits 1 – 6 and 4L (*nad1–6*, *nad4L*), and apocytochrome b (*cob*), genes for large and small subunit rRNA (*rnl*, *rns*), and a complete set of tRNA genes for mitochondrial translation using minimally-derived genetic code (Figure 1). The gene order was similar to that of other Bubarida (Lavrov et al. 2023) except for *nad1* having moved downstream of *nad5* and some modifications to tRNA gene locations. In addition, we identified two ORFs (*orf1535* of length 4608 bp and *orf167* of length 504 bp) upstream of *nad2*. Three genes contained group II introns: *cox1*, *cox2*, and *rnl*.

**Fig 1.**
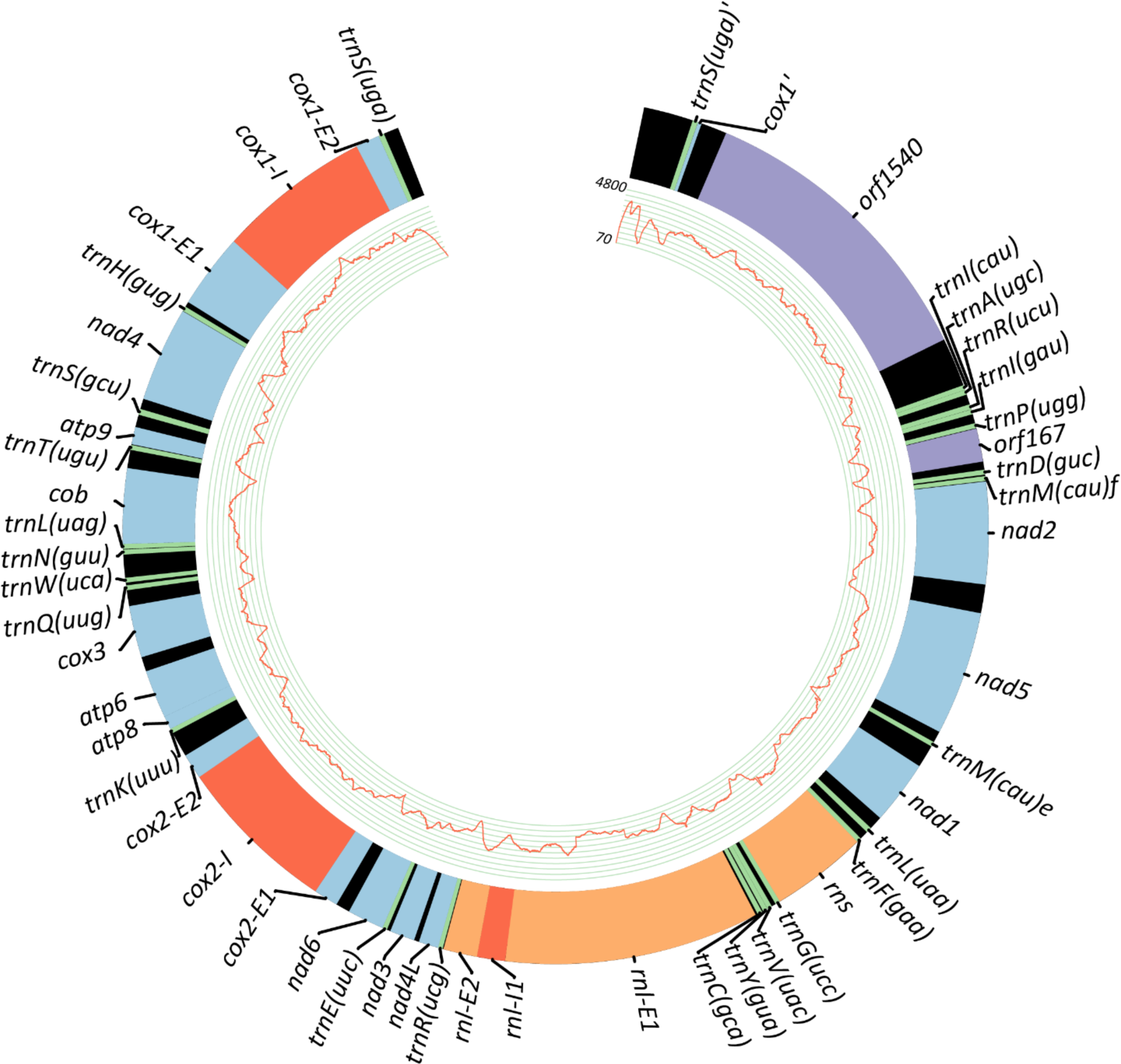
The gene order of the *Acanthella acuta* mitogenome. Coding regions are shown in blue, tRNAs in green, and rRNAs in orange. The *ORF1535* and *ORF167* are represented in purple, and group II introns are shown in red. Black bands represent intergenic regions. The line graph in the inner circles depicts the DNA coverage data for each position along the mitochromosome. *atp =* ATP synthase subunits, *cox =* cytochrome oxidase subunits, *nad =* NADH dehydrogenase subunits, *cob =* apocytochrome b, *rnl* = large subunit rRNA, *rns* = small subunit rRNA, E = exon, I = intron. tRNAs are represented by *trn* followed by their standard single-letter notation and anticodon in parentheses. trnM(cau)e = elongator methionine, trnM(cau)f = formyl methionine.

Noncoding regions ranging from 1 to 747 bp (average 142 bp) separate most genes, except for the following: *trnF, rns,* and *trnG* (no intergenic region); *trnR* and *nad4L* (no intergenic region); *atp6* and *atp8* (one nucleotide overlap).

### Novel features of A. acuta mitogenomes

When Lavrov et al. (2023) sequenced the *A. acuta* mitogenome, they were unable to PCR amplify a region upstream of *nad5* and downstream of *cox1.* Using total DNA Illumina sequencing we were able to reconstruct this region, revealing the presence of two large ORFs (*orf1535* and *orf167*, discussed below), *trnS*, an inverted copy *trnS (trnS’)* as well as a partial, inverted copy of *cox1* (*cox1’*). However, while assembling the mtDNA, no paired reads were found that would map beyond the terminal sequences (i.e., upstream of *orf1535* and downstream of *trnS*). We were also unable to PCR amplify across this region. These observations strongly suggest that the *A. acuta* mitogenome is organized as a single, linear molecule, as opposed to circular or multipartite.

Among the above-mentioned traits of the *A. acuta* mitogenome are two features that have previously been associated with linear mtDNA organization in some clades: 1) the presence of an ORF containing a DNA polymerase domain (*orf1535*), and 2) repetitive, inverted sequences at what appear to be the ends of the mitochromosome (ITRs).

*Orf1535* is 4,608 bp in length, located upstream of *nad2* and downstream of the ITR at one end of the chromosome, and is flanked by noncoding sequences of 453 bp and 747 bp in length up and downstream, respectively (Figure 1). Using InterPro scan (Paysan-Lafosse et al. 2023), we identified two different polymerase domains in *orf1535*, a DNA-directed DNA-polymerase (DNApol) domain and a DNA-directed RNA-polymerase (RNApol) domain. Maximum likelihood (ML) phylogenetic analysis of each domain separately showed that DNApol is most closely related to a putative DNA-directed DNA-polymerase from the demosponge *Amphimedon queenslandica* (A0A1X7TG59), with which it shares 15.56% amino-acid identity (Figure 2a). The RNApol domain grouped with mitochondrial DNA-directed RNA-polymerases from *Podospora* and *Neurospora* fungi (Q01521, P33541), with which it shared 15.51% and 18.24% amino acid identity, respectively (Figure 2b). Searching the IntroPro database for other proteins that harbored a similar gene fusion returned two hits: apricot (*Prunus armeniaca*, A0A6J5V), and quinoa (*Chenopodium quinoa*, A0A803N900). However, in both of these examples, the organization of the two domains within the protein was different from that seen in *A. acuta*, in that the proteins found in the plants had the RNApol domain upstream of DNApol. We were unable to find any other proteins on the InterPro database that have both these domains fused.

**Fig 2.**
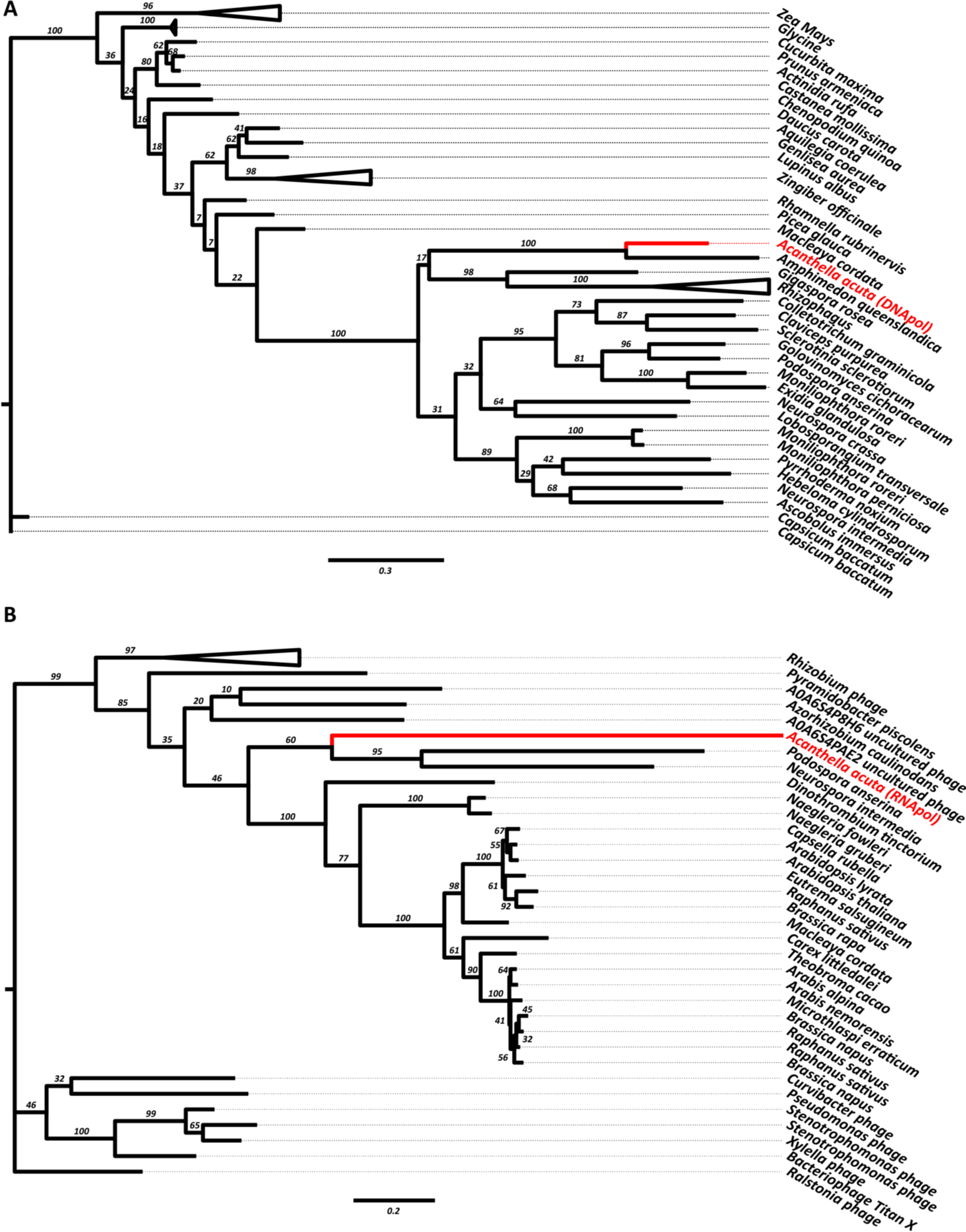
Maximum likelihood analysis of A) DNApol and B) RNApol domains of *orf1535* within the *A. acuta* (red) mitogenome. DNApol is most closely related to a gene found within the mitogenome of the demosponge *Amphimedon queenslandica*. RNApol is most closely related to a gene found within the mitochondrial genome of the fungi *Podospora anserina* and *Neurospora intermedia*. Bootstrap values of 200 replicates are shown.

In addition to *orf1535,* we identified another ORF, *orf167,* which was 504 bp in length and located between *trnP(ugg)* and *trnD(guc)*. This sequence does not return any hits when searched on the NCBI and UniProt BLAST tools and does not harbor any discernable protein domains.

We identified nearly identical but inverted sequences of length 371 bp at the terminal regions of the assembled *A. actua* mitogenome. These ITRs contain a part (55 bp) of *cox1* (*cox1’*) and a complete copy of *trnS* (*trnS’*), as well as noncoding segments. At the *orf1535* end of the chromosome, we asseebmed an additional sequence of 525 bp at the end of the molecule. The reverse complement of this sequence was not assembled on the opposite end of the chromosome, and its presence could not be confirmed through PCR. amplification. The partial sequence of *cox1* found in the ITR at the *orf1535* end is not duplicated on the other end, rather, the ITR on this end is contiguous with the complete *cox1* gene.

Three group II introns were identified in the *A. acuta* mtDNA: within *cox1* (2346 bp long, at position 1141), *cox2* (2426 bp long, at position 379), and *rnl* (431 bp). The *cox1* and *cox2* introns harbored an ORF encoding a reverse transcriptase domain. ML phylogenetic reconstruction for the *A. acuta cox1* intronic ORF grouped it with ORFs found within intron 1141 in four other sponges, where it was most closely related to that in *Phakellia ventilabrum*. The intronic ORFs from sponges were most closely related to those of red algae (Figure 3). Phylogenetic reconstruction for the *cox2* intronic ORF placed it with the green algae *Coleochaete* and *Nitella* (Figure 4). For the intron within *rnl*, the NCBI BLAST returned only six hits, all from placozoan *rnl*. These six sequences along with one other found within a dictyonellid sponge species, reported by Lavrov et al. (2023), were used for ML phylogenetic reconstruction. In the resulting phylogenetic tree, the *rnl* intron in *A. acuta* clustered most closely with the dictyonellid sponge intron (Figure 5).

**Fig 3.**
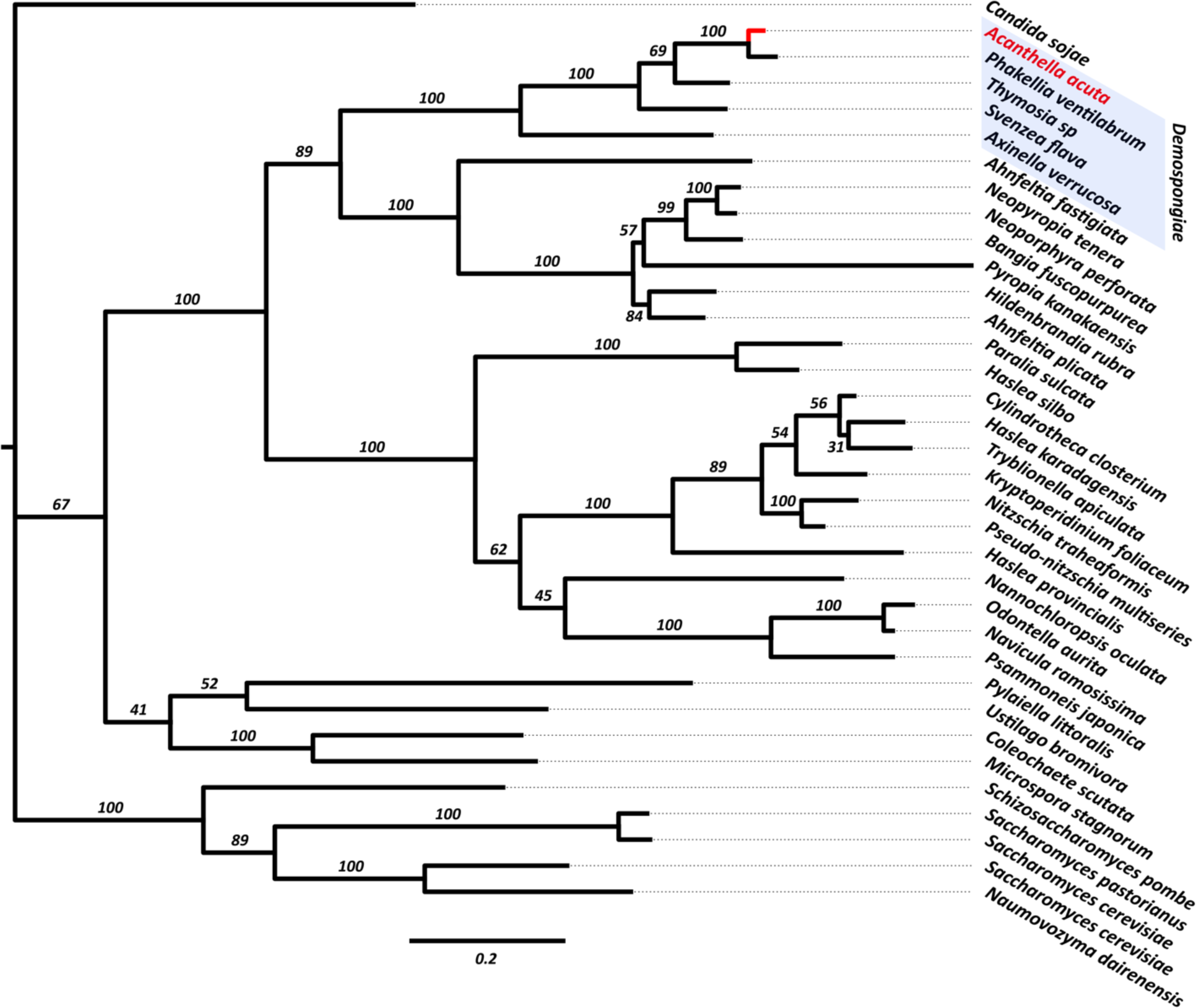
Maximum likelihood analysis of the *cox1* intron in the *A. acuta* (red) mtDNA. Demosponges are highlighted with a blue shaded box. Bootstrap values of 200 replicates are shown.

**Fig 4.**
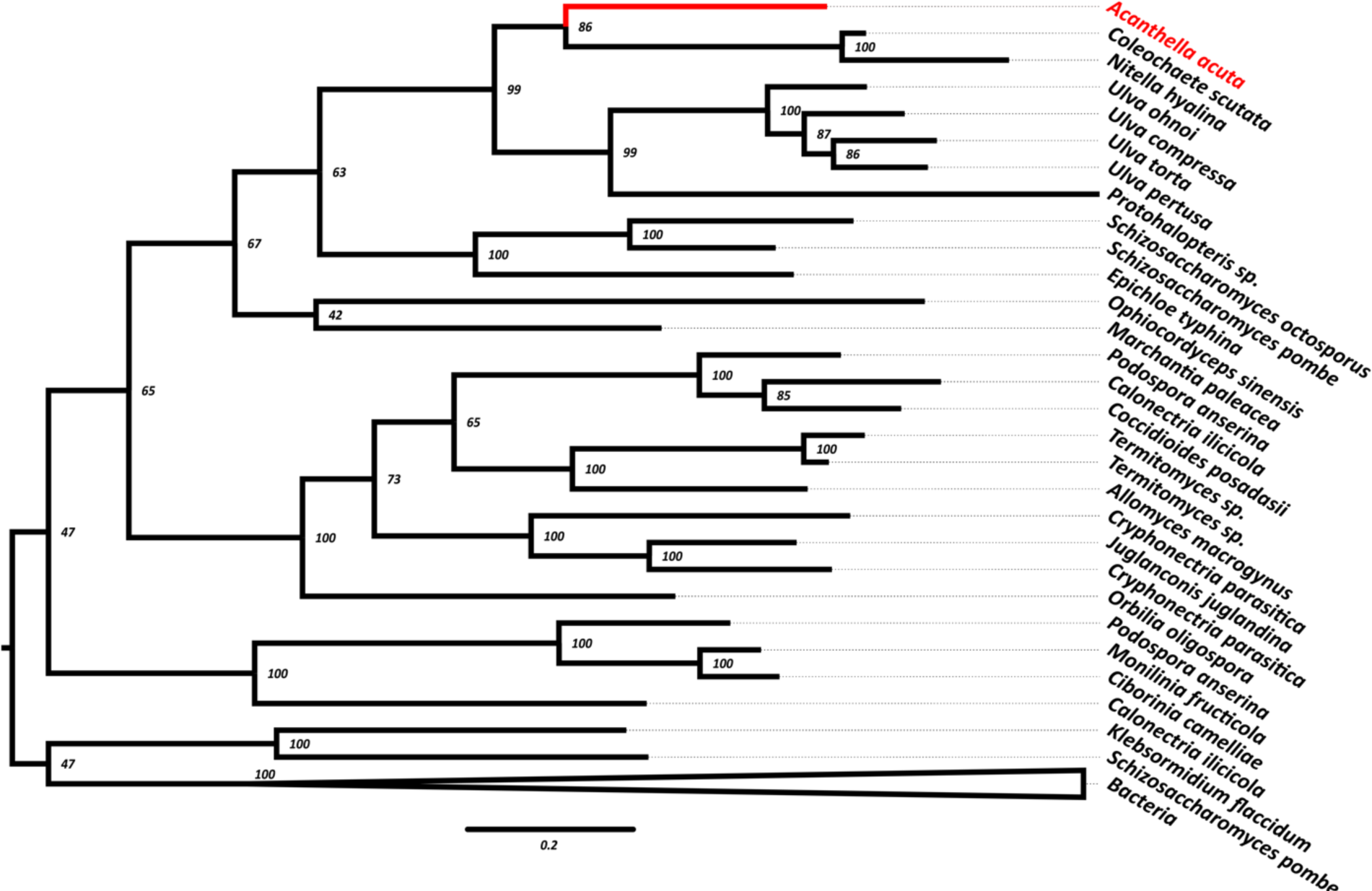
Maximum likelihood analysis of the *cox2* intron in the *A. acuta* (red) mtDNA. Bootstrap values of 200 replicates are shown.

**Fig 5.**
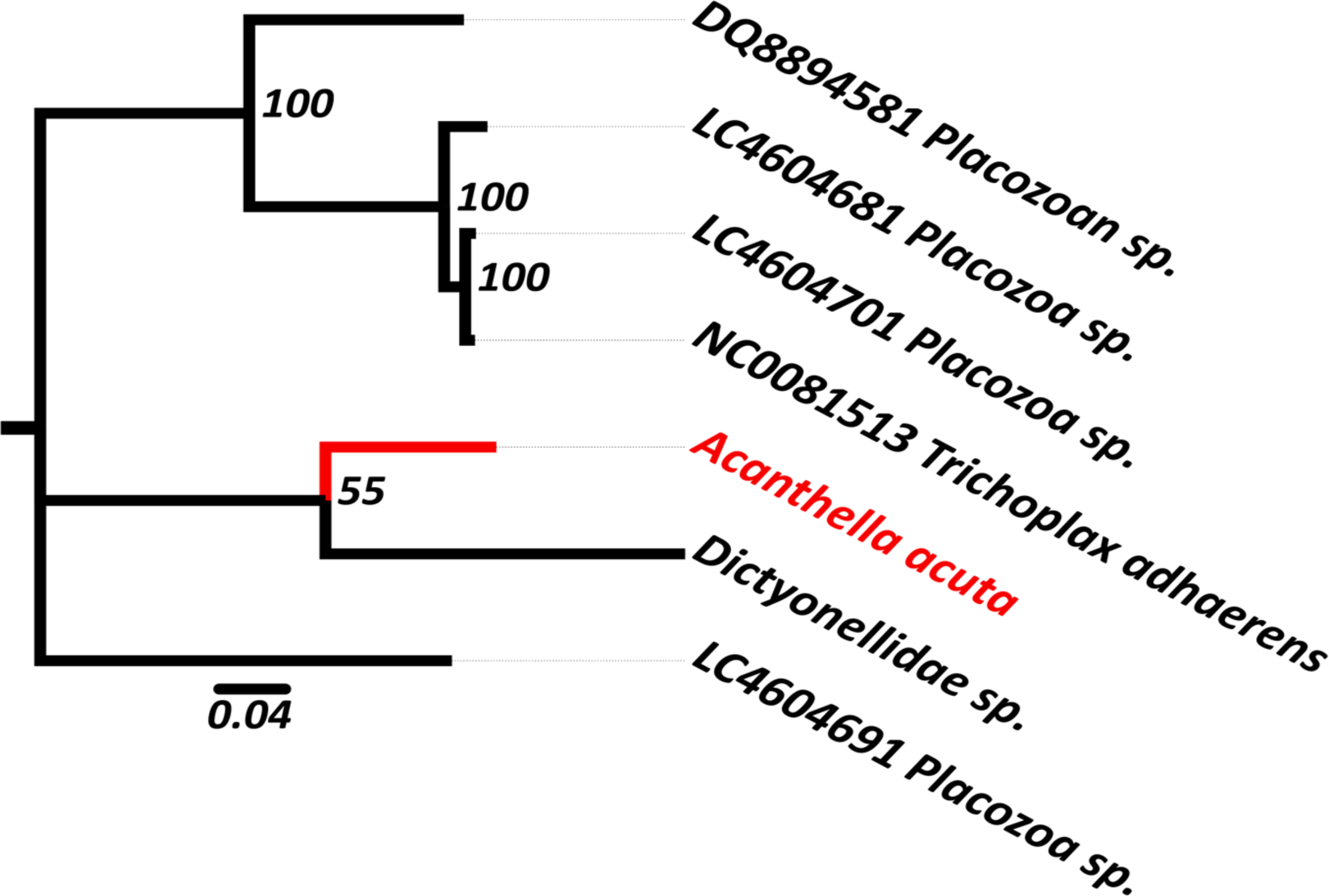
Maximum likelihood analysis of the *rnl* intron in the *A. acuta* (red) mtDNA. Bootstrap values of 200 replicates are shown.

We identified 21 short sequences that repeat multiple times across the mitogenome, both in forward (18 repeats) and reverse orientation (25 repeats). The longest of these, a sequence of 119 bp termed Short Repeat (SR) appears once in forward complement, and twice in reverse complement (SR’) and can fold into a structure containing four stem-loops. Most of the other sequences appear to be fragmented copies of SR, comprising complete or partial stem-loops. Figure 6A shows the alignment of the repeats against SR, and figurre 6B shows the folded structure of SR with how a subset of the other repeats relate to it. The lengths and number of instances for each of these sequences are given in Table 2

**Fig 6.**
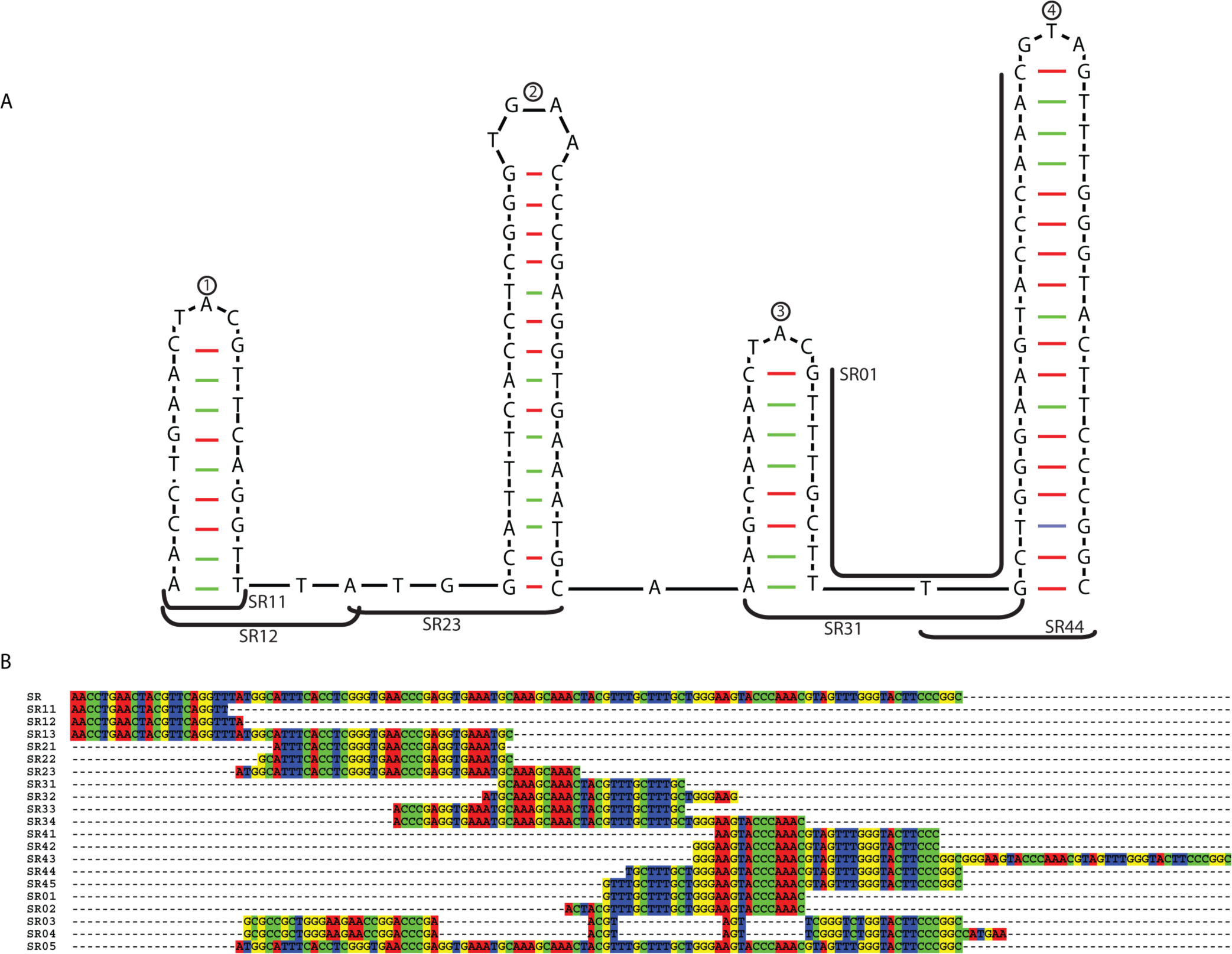
A) The nucleotide sequence of SR folded to form 4 stem-loop structures, labeled by the encircled numbers atop each. The locations of a subset of the other short repeat sequences are shown with labeled black lines. **B)** The alignment of all the short repeat sequences against SR. These sequences are named to represent which stem-loop in SR they align with (first digit), as well as their relative lengths (second digit) (therefore SR11 is the shortest sequence that aligns with the first stem-loop in SR, while SR23 is the longest sequence that aligns with the second stem-loop in SR. Sequences that do not align well with any stem-loop are termed SR0#).

**Table 2.** data on the 21 different Short Repeat sequences identified in the mtDNA.

| Name | Length | #forward repeats | #RC repeats | SR Loop |
| --- | --- | --- | --- | --- |
| SR | 119 | 1 | 2 | - |
| SR01 | 27 | 1 | 1 | - |
| SR02 | 32 | 0 | 1 | - |
| SR03 | 54 | 0 | 1 | - |
| SR04 | 60 | 1 | 1 | - |
| SR05 | 97 | 0 | 1 | 2, 3, 4 |
| SR11 | 21 | 3 | 5 | 1 |
| **SR12** | 23 | 3 | 2 | 1 |
| **SR13** | 59 | 0 | 2 | 1 |
| **SR21** | 31 | 1 | 0 | 2 |
| **SR22** | 34 | 2 | 1 | 2 |
| **SR23** | 46 | 1 | 2 | 2 |
| **SR31** | 25 | 1 | 0 | 3 |
| **SR32** | 34 | 0 | 1 | 3 |
| **SR33** | 39 | 1 | 0 | 3 |
| **SR34** | 55 | 0 | 1 | 3 |
| **SR41** | 30 | 0 | 1 | 4 |
| **SR42** | 33 | 1 | 0 | 4 |
| **SR43** | 36 | 0 | 2 | 4 |
| **SR44** | 45 | 1 | 0 | 4 |
| **SR45** | 48 | 1 | 0 | 4 |

## Discussion

### Linearization through plasmid integration

The mitochondrial DNA of *Acanthella acuta*, while typical in gene content and arrangement, is unusual in its size and genome organization. Its size – larger than any previously reported demosponge mtDNA – is attributed to the presence of two large ORFs (*orf1535, orf167*) and three introns. Most interestingly, the mitogenome of *A. acuta* is inferred to be linear with inverted repeats at the ends of the molecule, potentially acting as telomeres.

*A. acuta* is the first demosponge with linear mtDNA, and the fourth instance of mtDNA linearization within Metazoa (the others being Medusozoa, Calcarea, and some lineages of Isopoda). It likely acquired such an arrangement relatively recently, as inferred by the lack of linear DNA in other closely related species. The *A. acuta* mtDNA possesses an ORF containing a DNA-polymerase (DNApol) domain, another ORF of unknown function, and inverted terminal repeats (ITRs), much like what is assumed to be the ancestral mitochondrial organization in Medusozoa. It is, therefore, plausible that the mtDNA in *A. acuta* acquired its linear organization following the integration of a linear plasmid, as has been hypothesized for medusozoans (Kayal et al. 2012) and many species of fungi (Mouhamadou, Barroso, and Labare 2004).

Interestingly, *orf1535* contains an additional RNA-polymerase (RNApol) domain that has not been found in association with a DNA-polymerase domain in any animal documented to date. Furthermore, the two domains within *orf1535* do not share a common origin: the DNApol domain is more likely related to a nuclear gene found within the demosponge *Amphimedon queenslandica* while RNApol does not have any close relatives within the animal kingdom, clustering with fungi instead (Figure 2). This suggests that *orf1535* is likely the result of gene fusion (see below) of these domains. The correlation of this gene fusion with linear mtDNA architecture and whether these domains are functionally active is yet to be determined.

The ITRs of the *A. acuta* mitogenome contain a partial copy of *cox1* and a complete sequence for *trnS(uga).* These terminal repetitive regions may serve as telomeres for the linear mito-chromosome. Duplication of the end of the linear mitochondrial molecules has been reported in Hydrozoa (part of *cox1*) and Cubozoans (complete or partial coding sequences and RNA genes), among others (Kayal et al. 2012). Such duplications might be a result of a gene-conversion-based telomere maintenance system (Smith et al. 2012).

### Mitochondrial Introns

We identified group II introns within *cox1*, *cox2*, and *rnl* in the *A. acuta* mtDNA. Lavrov et al. (2023) previously reported the presence of a group II intron within *cox1* of numerous demosponges and suggested the presence of one within *cox2* of *A. acuta* as well. Introns, in general, are extremely rare in animal *cox2*, having only been reported in the bryozoan *Parantropora penelope* and the bivalve *Macoma balthica* (Lubośny et al. 2022). Specifically, group II introns in *cox2* have never been reported in animals before. Group II introns within *rnl* have only been reported in Placozoa (Burger et al. 2009) and one species of dictyonellid demosponge (Lavrov et al. 2023).

The introns found within *cox1* and *rnl* of the *A. acuta* mtDNA appear closely related to similar introns reported within other demosponges, suggesting that they have been inherited from a common ancestor. However, *A. acuta* is the only reported demosponge with an intron in *both cox1* and *rnl*.

Interestingly, the intron found within *cox2* of the *A. acuta* mitogenome does not appear similar to any reported in animals. Its close clustering with introns found within two charophyte algae suggests it might have invaded the *A. acuta* mitogenome through horizontal gene transfer. The *cox2* intron appears to be unique to *A. acuta* among demosponges and thus might be correlated to, or possibly acquired along with, the other unique features of this mitogenome (namely linear arrangement, *orf1535*, *orf167,* and ITRs; see below).

### Horizontal Gene Transfer and gene fusion

The *A. acuta* mitogenome possesses features that, given currently available data, are unique to this species within the animal kingdom: namely *orf1535* containing an RNA-polymerase domain, *orf167* and a group II intron within *cox2*. Phylogenetic reconstruction of the *cox2* intron and the individual RNApol domain (isolated by subdividing *orf1535*) suggests that these sequences are likely not of metazoan origin. In our reconstructions, the *cox2* intron was sister to sequences found in charophytes algae, while the RNApol domain was sister to sequences found in sordariales fungi. This suggests multiple horizontal gene transfer (HGT) events from various species to the *A. acuta* mitogenome. It is possible that the incorporation of these foreign sequences within the *A. acuta* mt-genome could correlate to its linear architecture. Alternatively, low sequence similarity scores of the closest matches of both *cox2* intron and RNApol domain may suggest that the common source of the HGT is just not known.

The *orf1535* is also unique in that it appears to be the result of a fusion between genes containing DNApol and RNApol domains. Our analysis suggests that these two domains do not share a common origin, and that, as previously stated, the RNApol domain is likely not of metazoan origin. We can think of three possibilities for how this fused ORF was acquired by *A. acuta*: 1) the two domains invaded the *A. acuta* mtDNA independently and fused after being incorporated, 2) the two domains underwent fusion in another organism’s genome and then invaded the *A. actua* mitogenome, or 3) since the DNApol domain appears to be related to a gene found within *Amphemadon queenslandica,* it is possible that this domain was not acquired through horizontal transfer, while the RNApol domain was, and the two domains then fused after the incorporation of RNApol into the mtDNA.

Given currently available data, scenario 2 seems to be the least probable situation, as we were unable to identify any other organism that contains a single ORF harboring both a DNA-polymerase and an RNA-polymerase domain in its mitogenome fused in the same order as *A. acuta*. Considering that close relatives of *A. acuta* do not appear to harbor a DNApol-containing ORF and that the similarity of DNApol to the *A. queenslandica* gene is low (15.56%), scenario 3 also looks unlikely. Therefore, in light of available data, scenario 1 appears to be the most likely situation, despite requiring two independent HGT events, followed by gene fusion.

While HGT might present a possible explanation for the presence of these unique intronic elements in the *A. acuta* mitogenome, it is more likely that this is merely an artifact of a limited representation of sponge genomes in publicly available databases. Indeed, when Huchon *et al*. (2015) reported the discovery of a group II intron in *cox1* of the demosponge *Axinella verrucosa*, their phylogenetic analysis led them to conclude that this intron had been horizontally transferred. Our initial phylogenetic reconstruction would have supported this inference, utilizing the available NCBI database. However, after the inclusion of Lavrov *et al.’s* recently gathered demosponge mitogenome data, it became apparent that *cox1* group II introns are widespread amongst demosponges, demonstrating that this intron has a deeper history within this group.

### Other Features

The *A. acuta* mitogenome contains multiple short repetitive sequences (SR#) that all appear to be identical to parts of the largest of these sequences, SR. It is possible that SR represents a remnant from an individual selfish element that invaded the A. acuta mitogenome at some point, and all the other SR# sequences are fragments of copies of SR that jumped to other parts of the mitogenome. It is also possible that the four stem-loops found in SR represent four individual elements that invaded the mitogenome and randomly converged to make the full SR sequence as they spread, although this situation seems unlikely. Whether the presence of these repetitive elements across the mitogenome relates to its linear architecture is unclear.

## Conclusion

*Acanthella acuta* is the first reported demosponge with linear mitochondrial DNA. The relatively recent origin of linear mtDNA in *A. acuta* may offer a unique opportunity to understand how linear mtDNA might emerge in animals, especially since it bears some similarities with linear mtDNA found in the ancient medusozoan lineage. The *A. acuta* mitogenome also possesses qualities that make it unique in the animal kingdom, such as a group II intron in *cox2*, and a never-before-seen fused ORF that could have resulted from multiple HGT events.

*A. acuta’s* unique mitogenomic organization also opens many new questions. For instance, what is the relationship between all the unusual features of the *A. acuta* mtDNA and its linear organization? Was the fusion of the two domains of *orf1535* concomitant with linearization? What is the origin of the *cox2* intron? Answering these questions would require further investigation into the mtDNA of Bubarida sponges, especially within the genus *Acanthella,* and would provide greater insight into the diversity and complexity of animal mitochondrial DNA.

## Methods

### DNA Extraction

A specimen of *Acanthella acuta* was collected by Manuel Maldonado (Centro de Estudios Avanzados de Blanes, Girona, Spain) and preserved in GuCl solution. Total DNA was extracted with a phenol-chloroform method modified from Saghai-Maroof et al. 1984 and used for library preparation.

Illumina True-Seq PCR free kit was used for the library preparation. Pair-end 100bp sequencing was done on Illumina NovaSeq 6000 at the Iowa State University DNA facility.

SPAdes assembler (v. 3.15.2) was used for contigs assembly and the mitochondrial contigs were identified using previously identified sequences from this species (Lavrov et al. 2019).

Genes and RNA sequences were annotated using the MFannot program by the University of Montreal (https://megasun.bch.umontreal.ca/apps/mfannot/). As recommended by the program, rRNA gene boundaries were adjusted based on comparisons with other nonbilaterian mitochondrial rRNA sequences (Altschul et al. 1990). The identities of the two *trnM(cau)* and *trnI(cau)* genes were established by sequence similarities with homologous genes in other demosponges.

The exact sequences at the ends of the linear molecule were identified by individually assembling both of the ends in isolation. For each end, a unique sequence of nucleotides near the end was used as a template: for the *orf1535* end, the sequence beginning at *orf1535* and extending to the stop codon of *nad2* was used, and for the *cox1* end, the sequence beginning at *nad4* and extending to the end of *cox1* was used. Illumina paired reads were isolated such that one read in the pair mapped to the assembled genome while its pair did not. These paired reads were then used in conjunction with the templates to assemble just the ends of the molecule utilizing the SPAdes assembler (version: 3.15.5).

### Phylogenetic Analysis

Phylogenetic analysis was conducted using sequences from the NCBI database for the *cox1* and *cox2* intronic ORFs (top 100 hits, E-value :: 2e-134) and the UniProt database for *orf1535* (top 50 hits). Additionally, for the *cox1* intron, three new sequences of *cox1* intronic ORFs were included from the demosponges *Phakellia ventilabrum, Svenzea flava,* and *Thymosia sp.* as described by Lavrov *et al*. (2023). Sequences were then filtered using CD-HIT with a similarity threshold of 95%. The filtered sequences were aligned using the MAFFT (v7.511) on auto settings. Highly variable sites in alignments were removed using GBlocks (-b5=a -p=n). Phylogenetic trees were constructed using RAxML-NG (v 1.0.2) with the WAG+G model for amino acid sequences (*orf1535* and the ORFs found within the *cox1* and *cox2* introns) and the GTR+G model for nucleotide sequences (*rnl* intron), and bootstrap values were calculated using 200 replicates.

## Acknowledgements

This work was supported by the National Science Foundation’s Assembling the Tree of Life program (DEB No. 0829783 to DVL) and by internal funds from Iowa State University. In addition, we would like to gratefully acknowledge Dr. Manuel Maldonado for sample contributed to this study.

## Data Availability

The WGS read files can be found under the NCBI SRA accession code SRR27117466, and the assembled and annotated mitochondrial genome can be found on the NCBI GenBank under the accession code OM729606.

